# Vernonia Amygdalina Del (Bitter Leaf) extract ameliorates isoniazid (INH) induced liver injury in Swiss Albino Mice

**DOI:** 10.1101/482091

**Authors:** Gebreselassie Addisu Tilaye, Muluken Fekadie Zerihun, Kasaw Adane Chuffa, Mahelet Arayaselassie, Daniel Seifu

## Abstract

Liver plays a central role in the metabolism of drugs. Drug clearance and transformation exposes liver to toxic injury. Antitubercular drugs have been found to be hepatotoxic and potentially lead to drug-induced liver injury. Isoniazid is one of the most hepatotoxic first line antitubercular drugs. Conventional drugs used in the treatment of liver disease are often inadequate and a search for supplementation or alternative drugs for the treatment of hepatic damage is indispensible. Therefore our study aims to investigate the hepatoprotective potential of *Vernonia Amygdalina Del* (bitter leaf) extract against Isoniazid-induced liver injury in Swiss Albino Mice. Treatment of Mice orally with *Vernonia Amygdalina Del* extract at dose of 250mg/kg and 375 mg/kg significantly lowered (P<0.05) the serum level of liver enzymes in Isoniazid pretreated mice. The hepatoptotective activity of the extract found to be comparable with the standard drug, Silymarin (100 mg/kg, P.o.). Moreover, treatment with the extract significantly alleviated Isoniazid induced hepatic injury as supported by the photomicrographs of liver section of mice. The data shows aqueous *Vernonia Amygdalina Del e*xtract has a very promising hepatoprotective potential against isoniazid-induced liver injury.

## Introduction

Liver is the largest organ of the body weighing approximately 1500g, located in the upper right quadrant of the abdomen, anterior to the right kidney and inferior to the diaphragm [1]. The organ is responsible for over 500 metabolic functions and maintenance of hemeostasis [2] synthesis of aminoacids, plasma proteins and many biochemicals, clotting fators, gluconeogenesis, and glycogenolysis, and urea production [3]. Liver can serve as a storage organ for several products like glycogen, fat, and fat soluble vitamins stored within the liver parenchyma. It produces bile which is secreted into the lumen to assist fat digestion process and purifying the blood [4]. Excess production and/or accumulation of toxic chemicals hinder the production of the bile leading to body’s inability to flush out chemicals through waste. Smooth endoplasmic reticulum of the liver is the principal organelle serves as a “metabolic clearing house” for endogenous chemicals like cholesterol, steroid hormones, fatty acids and proteins and exogenous chemicals such as drug and alcohol.

Drug clearance and transformation roles of the liver exposes the organ to toxic injury [4]. Hepatotoxicity can be caused by over doses of certain medicinal drugs, industrial chemicals, herbal remedies, and even dietary supplements [5]. Drug-induced hepatotoxicity is the most common cause of acute liver failure in many countries [6]. Certain drugs might even cause such injuries within therapeutic range and the severity of liver injury greatly increased if drug is continued after the onset of symptoms.

Antitubercular drugs have been found to be potentially hepatotoxic and often lead to liver injury. Anti-tuberculosis drug-induced hepatotoxicity (ATDH) is main cause of treatment interruption and change in treatment regimen during tuberculosis treatment process. Isoniazid, rifampicin and pyrazinamide are well known first-line anti-tuberculosis drugs [7]. Isoniazid has been widely used for treatment of tuberculosis. However, if the drug is consumed in overdose or for a long period of time susceptible individuals might suffer sever hepatotoxicity. Its effect may range from mild increase in serum liver enzyme activity to sever forms of hepatocellular necrosis or intrahepatic cholestasis [8]. Mild and transient serum enzyme increases occur in 10–20% of the patients taking the drug and severe hepatotoxicity occurs in about 1–3% of patients [9]. Isoniazid is the most hepatotoxic antitubercular drug [7, 8]. The drug received a black box warning from the US Food and Drug Administration (FDA) due to its high incidence of adverse drug reactions in resulting hepatocyte injury [10]. Various metabolites of isoniazid have been suggested as being hepatotoxic, including hydrazine, monoacetyl hydrazine, acetylisoniazid and isonicotinic acid [11]. Administration of acetylhydrazine or acetylisoniazid in rats leads to the production of reactive alkylating species that covalently bind to liver proteins, causing hepatocyte injury [12]. In another study, Isoniazid induced toxicity resulted a significant elevation in the level of Liver enzymes, alanine aminotransferase (ALT), aspartate aminotransferase (AST) and alkaline phosphatase (ALP) [13]. Therefore in this study we used Isoniazid to induce hepatotoxicity in Swiss Albino Mice and the doses of Isoniazid used to induce hepatotoxicity was adopted from Chen *et al.*, 2011 [14].

Although Isoniazid (INH) is known to be potentially hepatotoxic, because of its efficacy the drug remains a mainstay for the treatment of tuberculosis [15]. Conventional drugs used in the treatment of liver disease are often inadequate. It is therefore search for supplementation/ alternative drugs for the treatment of hepatic damage caused by anti-tubercular drugs.

## Materials and methods

A total of 200g shade dried leaves were powdered in an electrical grinder and soaked in 1.8 litres of distilled water. The filtrate was frozen and lyophilized to dryness. After drying, a dark brown extract of *Vernonia Amygdalina* was stored at 4^0^C until tested in the animals. The animals were assigned to one of six groups, each group consisting of six mice. The control mice were orally injected with sterile saline (1 mL/kg). Liver injury was induced by daily dose of isoniazid (75 mg/kg, P.o.) for two weeks as manifested by statistically significant elevation in serum ALT, AST, ALP and bilirubin level. An obvious death of hepatocytes, accompanied by mild necrosis and infiltration, was observed in liver of mice treated with isoniazid. Positive control mice received INH at 75mg/kg p.o. daily PLUS silymarin orally at 100mg/kg daily. The treatment groups received a daily dose of (250 mg/kg and 375 mg/kg) *Vernonia Amygdalina* extract (P.o) soon after (75mg/kgP.o) isoniazid was administered.

### Study setting

The study was conducted at Addis Ababa University, Department of Biochemistry.

### Plant collection

*Veronia Amygdalina Del* leaves were collected in Addis Ababa Ethiopia in September 2013. The specific plant species were confirmed by taxonomist working at Addis Ababa University (AAU) National Herbarium. Voucher specimens were dried and deposited (Voucher number 001, September, 2013) at AAU, Ethiopia.

### Extraction procedure

Leaf chemical extraction of *Vernonia Amygdalina Del* was done at Addis Ababa University, Department of Pharmacology laboratory. Fully green *Vernonia Amygdalina Del* leaves were collected from Lideta subcity, Addis Ababa, Ethiopia. Leaves were washed with distilled water in dust protected laboratory room. Two hundred gram shade dried leaves were powdered in an electrical grinder and soaked in 1.8 litres of distilled water. The well mixed mixture was kept in the laboratory for 24hrs before filtering. The filtrated was frozen and lyophilized into dryness. Finally dark brown 18g (9%) of the original specimen extract of *Vernonia Amygdalina* was obtained and stored at 4°C until tested in animal model.

### Ethical approvals

Ethical clearance was obtained with protocol No. 010/13 from research and ethics committee of the Department of Biochemistry, Addis Ababa University.

### Preparation of animals

Thirty-six female SAM weighing 22g to 30g having three months of age were obtained from the department of Pharmacology, Addis Ababa University. Random assignment of mice to six groups of each six mice per cage was done. Food and water was supplied (standard food pellets) adlibitum, ambient temperature 21°C and 12hrs light/dark cycle. The animals were allowed to acclimatize for 2 weeks before the experiment. Cages were cleaned daily, food and water was changed daily. All animals were inspected and/or observed for food and water intake during the time of treatment.

### Study design and experimental design

Randomized controlled experimental design was employed to undertake this study on animal model. Experiments were carried out according to the guidelines for care and use of experimental animals. Each female SAM was randomly assigned to any of the six experimental groups.

✓ The Group I, the control (non-treated) group of six mice were orally injected with sterile saline (1 mL/kg) and were not treated with isoniazid and *Vernonia Amygdalina Del* extract.
✓ Group II (six mice) received *Vernonia Amygdalina Del* extract alone orally at a dose of 375 mg/kg [16] once daily.
✓ Group III (six mice) received only INH, at a dose of 75mg/kg p.o. once daily
✓ Group IV (six mice) received INH at a dose of 75mg/kg p.o daily PLUS *Vernonia Amygdalina Del* extract orally at a dose of 250 mg/kg [16] daily.
✓ Group V (six mice) received INH at 75mg/kg p.o daily PLUS *Vernonia Amygdalina Del* extract orally at 375 mg/kg [16] daily.
✓ Group VI (six mice) received INH at 75mg/kg p.o. daily PLUS silymarin orally at 100mg/kg daily.

### Drugs and chemicals

The first-line anti-tuberculosis (anti-TB) or isoniazid (INH) drug was obtained from the department of pharmacy, Addis Ababa University. The leaf extract was dissolved in sterile distilled water and analytical grade reagents. Isoniazid (INH) causes significant liver injury on mice more than it results in rats and human with slow acetylators have comparable affects where the former with increased risks of hepatotoxicity [17].

### Isoniazid model for evaluation of antihepatotoxic activity

The isoniazid model was used for scheduling the regimen of dose 75mg/kg drug diluted in sterile distilled water once a day to induce liver damage [14].

### Anaesthesia and blood collection procedure

Animals were anesthetized with diethyl ether via cotton inhalation before cardiac puncture. 1mL of blood specimen was collected from each mouse using 3mL of syringe after two weeks of daily treatment. The blood specimen after left to clot for 30 minute serum was separated by centrifuging at 4500 r.p.m for 3 minutes. Five hundred micro liter of non-hemolized serum was separated and stored in a deep-freezer at −40°C until blood chemistry analysis of liver function enzymes (ALT and AST), cholestatic markers (ALP and total bilirubin) and serum total protein.

### Biochemical assessments

The synthesis of several plasma proteins and detoxification and excretion of bilirubin is regulated by the liver. Measurement of bilirubin, lipid, lipoprotein, and some other important proteins are regulated as indicator of liver function. Moreover; determination of serum liver enzyme activities of ALT, AST, and ALP is considered as good marker of hepatic injury or hepatocellular integrity.

### Serum biomarkers for liver function tests and total protein level

#### Alanine aminotransferase (ALT) assay

Alanine aminotransferase (ALT) or serum glutamate-pyruvate transferase (SGPT) is mainly expressed in the liver and its increased level is indicator of liver injury [18]. This has been used in assessing preclinical investigation of experimental drug formulation on rodents [19].

### Reaction principle

The reagent cuvette was incubated at a temperature of (37°C ± 0.5 °C) for the duration of the test, then 50 µl of serum sample was mixed with 500µl of the working reagent and incubated for 1 min at 37 °C. The absorbance was read at 340 nm exactly after 1, 2, and 3 minutes. The mean absorbance of change per minute was computed to calculate the activity of ALT. The conversion of NADH to NAD+ correlates with serum ALT activity, and which is determined by continuously monitoring the loss of NADH absorbance at 340 nm by spectrophotometer.

### Aspartate aminotransferase (AST) Assay

Aspartate transaminase (AST) or serum glutamate-oxaloacetate transaminase (SGOT) is an enzyme expressed in the liver. The enzyme was determined from serum samples based on the kinetic method. Evaluation of the level of AST indicates the magnitude of acute liver damage or liver injury [18] but not specific to liver only. However; it has been used to monitor hepatotoxic effects of experimental drugs in rodents [19]. The ratio of AST to ALT is sometimes helps in differentiating causes of liver damage. For instance, AST/ALT elevations instead of ALP elevations favour liver cell necrosis as a mechanism over cholaestasis. AST and ALT are both over 1000 IU/L, the differential can include acetaminophen toxicity, shock, or fulminant liver failure. When AST and ALT are greater than three times normal but not greater than 1000 IU/L, the differential can include alcohol toxicity, viral hepatitis, drug-induced, liver cancer, sepsis, Wilson’s disease, post-transplant rejection of liver, autoimmune hepatitis, and (nonalcoholic) steatohepatitis [20].

### Reaction principle

The reagent cuvette was incubated at (37 °C ± 0.5°C) during the testing procedure. 50 µl of serum sample was mixed with 500µl of the working reagent and incubated for 1min at 37 °C. The absorbance was read at 340 nm exactly after 1, 2, and 3 minutes. The mean absorbance change per minute was calculated and the concentrations will be obtained by using the formula. The conversion of NADH to NAD+ correlates with serum AST activity, and is determined by continuously monitoring the loss of NADH absorbance at 340 nm by spectrophotometer.

## Cholestatic indices

### Alkaline phosphatase (ALP) assay

Alkaline phosphatase (ALP) catalyzes the hydrolysis of phosphate esters in an alkaline environment, resulting in the formation of an organic radical and inorganic phosphate. In mammals, this enzyme is found mainly in the liver and bones. Marked increase in serum ALP levels has been associated with malignant biliary obstruction, primary biliary cirrhosis, hepatic lymphoma and sarcoidosis, and several bone diseases [18]. The enezyme hydrolyze p-Nitrophenylphosphate (pNPP) releasing p-Nitrophenol (pNP) and Phosphate with a yellow colored product. The rate of pNP (quinonid form) release phosphatase activity which is determined by continuous monitoring of the increase in absorbance at 405 nm.

Reaction principle

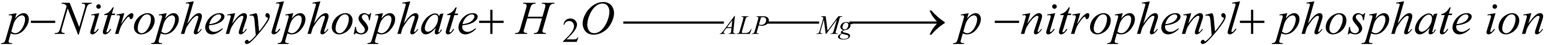

### Procedure

The reagent cuvette was incubated at (37 °C ± 0.5 °C) for the duration of the test. Then 20 µl of serum sample was mixed with 500 µl of the working reagent and incubated for 1min at 37 °C. The absorbance was read at 405 nm exactly after 1, 2, and 3 minutes and from the reading the mean absorbance change per minute was calculated. The rate of the reaction is directly proportional to the alkaline phosphatase activity. Finally the activity of alkaline phosphatase in the sample was calculated from the mean absorbance change per minute.

The level of ALT, AST, ALP and total protein were analyzed using Auto lab 18 Analyzer (fully automated chemistry analyzer, Italy).

## Total bilirubin assay (Jendrassik-Grof) principle

### Principle

Total bilirubin was measured using Jendrassik-Grof reaction method in the presence of accelerator using diazonium salt blue-green coloured mixture at PH of 13. The intensity of the colour is read at 600 nm and it is directly proportional to the concentration of total bilirubin.

### The Reaction

Bilirubin + diazotized sulfanilic acid ? azobilirubin

The alkaline reaction produces a more intense colour than the equivalent reaction run at a neutral pH [18]. Total bilirubin coupled with a diazoniumsulphanic acid yield the corresponding azobilirubin in the presence of methanol or urea accelerators. The values measured in the absence of solubilizers (for example, methanol) is called direct bilirubin where as in their presence it is total bilirubin (Table 1).

**Table 1:** Assay procedures of bilirubin determination.

| Reagents | Sample blank(μL) | Sample test(μL) |
| --- | --- | --- |
| Serum sample | 100 | 100 |
| Distilled water | - | - |
| Methanol | 500 | 500 |
| 1.5% HCl | 100 | - |
| Diazo (diazotized sulfanilic acid) | - | 100 |

### Procedure

Sample blank and test specimen are incubated at 37°C for 5 minutes and then read by the semi-automated 5010 photometer chemistry analyzer.

## Statistical Tests

The magnitude of the effect of the intervention among experimental groups of animals was done by using SPSS statistical software package version (version 16) to evaluated using independent t-test and p-value less than 0.05 was considered as statistically significant.

## Variables

### Dependent variables

- Concentration of liver enzymes
- Total bilirubin in serum
- Liver/body weight %
- Histopathological changes

### Independent variables

- *Vernonia Amygdalina Del* extract

### Tissue examination for histopathology

Histological slides from each of six groups were examined by pathologist for any liver pathological change.

### Collection preparation and staining of tissue sample

Finally animals were sacrificed and tissue specimen from the right lobe was taken and transferred into 10% buffered neutral formalin [21] fixes, inhibits decay, autolysis or degeneration of tissues. Fixed tissues were washed in running tap water for 8 hrs to allow paraffin wax easy infiltration into the tissue [22].

Tissue specimen were dehydrated by immersing it an increasing concentration of 70%, 80%, 95%, 100% [23]. Clearing was done by immersing tissues in two steps of xylene each 1 hr to remove ethanol and replace it with paraffin miscible fluids [22]. Two changes of paraffin wax each 1 ½ 56°C (52-64°C) enables infiltration [23].

Paraffin embedded tissue blocks were moulded square using metallic plate and electro-thermal wax dispenser. The blocks were the labelled and sealed in plastic bags with the examining surface downward prior to sectioning were placed in a refrigerator until sectioned [23] which gives too small and/or delicate firm tissues surrounded with wax [22]. The tissue blocks were put in the rotary microtome and sectioning of tissue blocks produces sections of 5 μm. The section ribbons were carefully picked using blunted forceps and let to float in a water-bath adjusted at 40°C (slightly below the melting point of wax) to unfolds the sections. Unfolded sections were transferred onto clean microscopic glass slides pre-incubated in an oven at 56 °C for 20-30 minutes for better drying and adhesion. At this stage, the sections were ready for staining [23] using Clopton’s formula [24].

The paraffin was removed from the tissue sections by immersing in a series of descending order alcoholic concentration. Distilled water removes xylene and tissue hydration is attained. Hydrated sections were then immersed in hematoxylin for 3-5 minutes with an eosin counter stain and agitated with acid alcohol to prevent over-staining. Sections were immersed in a mixture of sodium bicarbonate, ethanol, and distilled water or tap water to give blue colour to the nucleus. Finally, it was immersed in 95% alcohol and eosin to give pink colour to the cytoplasm [23, 24].

Tissue sections were then dehydrated in 95% alcohol, cleared in xylene, and mounted by adding a drop of DPX (Dibutyl phthalate in xylene) mounting medium which cover the microscopic glass and increase the refractive index of the tissue under light microscope. This prevents bubble formation between the tissue and the cover glass [22].

## Results and discussion

### Vernonia Amygdalina Del extract restored serum level of liver enzymes in isoniazid (INH) challenged mice

Treatment with aqueous *Vernonia Amygdalina Del* extract restored serum level of liver enzymes near to normal in isoniazid challenged mice. *Vernonia Amygdalina Del* treatment at a dose of 250 mg/kg (group-4) showed significant reduction (P<0.05) of ALT level in isoniazid pretreated mice compared to only isoniazid treated group (group-3). Hepatotoxicity was induced by daily administration of anti-tubercular drug (isoniazid) (75mg/kg P.o.) for 15 days as confirmed by significant elevation of serum level of liver enzymes such as ALT (P<0.01), ALP (P<0.01) and AST (P<0.05) levels compared to the sterile saline (1mL/kg) treated control group (Table 2). At the time of hepatic injury, these enzymes leak out from liver into the systemic circulation due to liver tissue damage. An obvious death of hepatocytes, accompanied by mild necrosis and infiltration, was also observed in the liver of mice treated with only isoniazid. However, no significant difference in the level of liver enzymes was observed between the sterile saline (1mL/kg) treated group and only *Vernonia Amygdalina Del* extract treated group (375 mg/kg) indicating the extract itself did not affect the normal physiology of the liver. Moreover, treatment at a higher dose (375 mg/kg) group-5 lowered the level of liver enzymes near to normal (P<0.01) in isoniazid pretreated mice compared to INH only treated group. Treatment with silymarin (group-6) also showed significant reduction (P<0.01) in the level of liver enzymes in isoniazid pretreated mice compared to INH only treated group.

**Table 2:** Effect of *Vernonia Amygdalina* on liver enzyme parameters in isoniazid treated mice.

| Groups | ALT (U/I) | AST (U/I) | ALP (U/I ) |
| --- | --- | --- | --- |
| Group-1 sterile saline (1mL/kg) | 29.80 ±4.45 | 114.0 ±6.52 | 75.96±6.58 |
| Group-2 (375 mg/kg) *Vernonia Amygdalina* extract (P.o.) | 30.88 ± 3.81 | 115.0±4.47 | 76.64± 4.57 |
| Group-3 Isoniazid (75 mg/kg, P.o.) | 39.42 ± 4.19<sup>a</sup> | 125.3 ±7.69 | 90.02 ± 5.80<sup>a</sup> |
| Group-4 Isoniazid (75 mg/kg, P.o.) + (250 mg/kg) *Vernonia Amygdalina* extract (P.o.) | 32.74 ± 2.43<sup>b</sup> | 117.5 ±1.78 | 80.24± 6.56<sup>b</sup> |
| Group-5 Isoniazid (75 mg/kg, P.o.) + (375 mg/kg) *Vernonia Amygdalina* extract (P.o.) | 29.34± 2.78<sup>c</sup> | 114.0 ±6.49<sup>b</sup> | 79.12± 4.58<sup>b</sup> |
| Group-6 Isoniazid (75 mg/kg, P.o.) + silymarin orally at 100mg/kg daily | 28.22± 2.75<sup>c</sup> | 114.4 ±5.15<sup>b</sup> | 79.14± 4.70<sup>b</sup> |

<sup>a</sup>:p<0.01 vs group-1; <sup>b</sup>:p<0.05 vs group-3; <sup>c</sup>:p<0.01vs group-3

### *Vernonia Amygdalina* extract restored serum total bilirubin and liver/body weight %

Treatment with *Vernonia Amygdalina Del* extract significantly reduce serum total bilirubin in Isoniazid pretreated mice (P<0.05) in isoniazid PLUS *Vernonia Amygdalina Del* extract at doses of 250 mg/kg) and (P<0.01) in isoniazid PLUS *Vernonia Amygdalina Del* extract at doses of 375 mg/kg)) compared to Isoniazid only treated mice (Table 3). Since, serum level of bilirubin reflects the liver’s ability to take up, process, and secrete bilirubin into the bile the result of this study indicates the regeneration of the liver to perform its physiological function. However, there was no significant difference in the level of total bilirubin between the sterile saline (1mL/kg) treated control and only *Vernonia Amygdalina Del* extract (375 mg/kg) treated group. The data indicates that *Vernonia Amygdalina Del* extract has no significant undesired effect on hepatocellular integrity on its own. Moreover, significant decrease (P<0.01) in the level of total bilirubin was observed in silymarin (100mg/kg) treated group compared to only Isoniazid treated group. Moreover, the data on Table 3 indicated that *Vernonia Amygdalinadel* treatment resulted in a significant decrease (p<0.01) in liver/body weight % in isoniazid-pretreated group compared to only Isoniazid treated group. This is probably due to the healing of liver to perform all its physiological functions including burn the fat it had accumulated inside its cells.

**Table 3:**
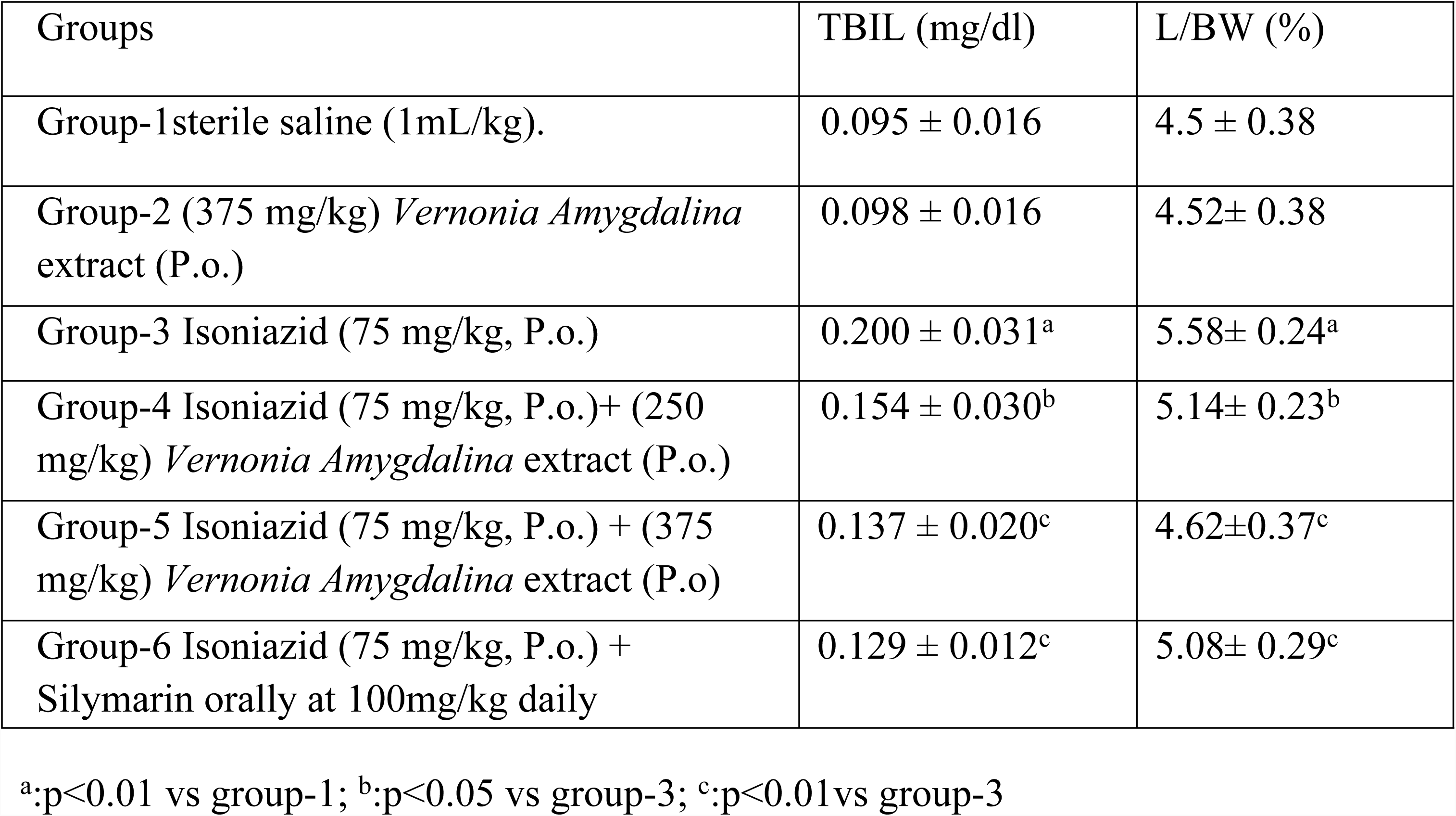
Effect of *Vernonia Amygdalina Del* on total bilirubin and liver/body weight % in isoniazid treated mice.

### *Vernonia Amygdalina Del* regenerated the liver architecture in isoniazid-pretreated mice

Microscopic examination of the liver sections of mice showed visible difference in the liver architecture between the controls and the treatment groups. As we can see from the microscopic slide the sterile saline (1mL/kg) treated control group shown normal liver architecture, normal hepatocytes containing central vein and sinusoid and mice treated orally with the aqueous leaf extract of *Vernonia Amygdalina Del* at a dose of 375 mg/kg (group-2) showed no significant changes in their liver architecture compared to the control. In contrast, the liver histology of only Isoniazid treated group showed centrilobular necrosis with a mild lymphocytic infiltrate, a change in the shape of the central vein, size of hepatic sinusoids and small number of hepatocytes compared to the control.

However, treatment of mice orally with aqueous leaf extract of *Vernonia Amygdalina Del* significantly regenerated the liver architecture in isoniazid-pretreated mice (fig 1D) and a dose dependent difference in regenerative capacity was observed between group-4 (250 mg/kg) (fig 1D) and group-5 (375 mg/kg) (fig 1E). The silymarin treatment (fig 1F) also significantly regenerated the liver architecture in Isoniazid pretreated mice (isoniazid PLUS 100 mg/kg silymarin) compared to only Isoniazid treated group (fig 1C) even though there is minor infiltration. The result of *Vernonia Amygdalina Del* at a dose of 375 mg/kg was comparable to silimarin treated groups.

### Percentage yield of aqueous leaf extract of *Vernonia Amygdalina Del*

A total of 200 grams of shade dried *Vernonia Amygdalina Del* leaves were powdered in an electrical grinder and soaked in 1.8 liters of distilled water. The filtrate was frozen and lyophilized to dryness. After drying, a dark brown extract of *Vernonia Amygdalina Del* weighing 18 g was obtained.

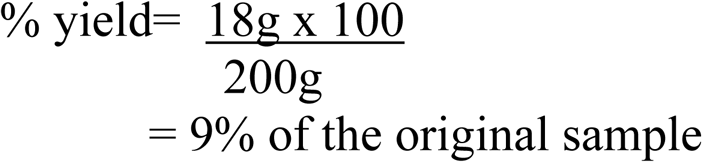

Treatment with *Vernonia Amygdalina Del* restored serum level of liver enzymes near to normal in isoniazid challenged mice. This study has further demonstrated the hepatoprotective potential of the plant via histopathological investigation. The finding of this study appears to validate the earlier observation of Iwalokun et al., 2006 that the terpenoid fraction of *Vernonia Amygdalina* leaf extract ameliorates acetaminophen induced hepatotoxicity in mice [25]. Furthermore, the treatment with *Vernonia Amygdalina Del* extract restored the level of bilirubin to near normal in isoniazid pretreated mice.

The treatment with *Vernonia Amydalina Del* also regenerated the isoniazid induced histopathological changes in the livers of mice. It is suggested that the hepatoprotective activity of *Vernonia Amygdalina Del* against isoniazid induced hepatotoxicity might be due to its property of inhibiting several isoforms of cytochrome P450 enzymes, and that it potentiates the antioxidant capacity of the liver, acts as a scavenger of oxygen free radicals, and inhibits the synthesis of proinflammatory cytokines. In addition to its hepatoprotective actions, *Vernonia Amydalina Del* has been shown to be effective in inhibiting tumor growth and promotion in several types of cancer.

Liver disease is a worldwide problem. Conventional drugs used in the treatment of liver diseases are sometimes inadequate and can have serious adverse effects. It is therefore necessary to search for alternative drugs for the treatment of liver diseases to replace currently used drugs of doubtful efficacy and safety. The results of the present study, along with the above facts, strongly suggest that aqueous leaf extract of *Vernonia Amygdalina Del* has hepatoprotective potential against Isoniazid induced liver injury, which can be attributed to the presence of terpenoids, alkaloids and flavonoids.

## Conclusions

Treatment with *Vernonia Amygdalina* restored serum level of liver enzymes near to normal in isoniazid challenged mice. This study has further demonstrated the hepatoprotective potential of the plant via histopathological investigation. The finding of this study appears to validate the earlier observation of Iwalokun *et al.*, 2006 that the terpenoid fraction of *Vernonia Amygdalina* leaf extract ameliorates acetaminophen induced hepatotoxicity in mice. Furthermore, the treatment of *Vernonia Amygdalina* restored the level of bilirubin to near normal in isoniazid pretreated mice. The treatment with *Vernonia Amydalina* also normalized the isoniazid induced histopathological changes in the livers of mice. It is suggested that the hepatoprotective activity of *Vernonia Amygdalina* against isoniazid induced hepatotoxicity might be due to its property of inhibiting several isoforms of cytochrome P450 enzymes, and that it potentiates the antioxidant capacity of the liver, acts as a scavenger of oxygen free radicals, and inhibits the synthesis of proinflammatory cytokines. In addition to its hepatoprotective actions, *Vernonia Amydalina* has been shown to be effective in inhibiting tumor growth and promotion in several types of cancer. Liver disease is a worldwide problem. Conventional drugs used in the treatment of liver diseases are sometimes inadequate and can have serious adverse effects. It is therefore necessary to search for alternative drugs for the treatment of liver diseases to replace currently used drugs of doubtful efficacy and safety. The results of the present study, along with the above facts, strongly suggest that aqueous leaf extract of *Vernonia Amygdalina* has hepatoprotective properties, which are mediated by antioxidant activity in mice in vivo.

There was a problem of developing an animal model of isoniazid-induced hepatotoxicity. Because of its short half-life, it was found that smaller, more frequent doses of isoniazid lead to greater hepatotoxicity than one large dose. But we managed this challenge by providing the mice with the minimum toxic dose of the drug. The other challenge was it is recommended to give the drug in food because isoniazid has a short half-life, and this provided a more consistent blood level than once-a-day oral gavages. This method of drug administration produced blood levels in mice that were comparable to the isoniazid in humans. However, the mice were not voluntary to eat the pellet mixed with the drug. So we managed the problem by giving the drug in a solution via oral gavage.

## Abbreviations

AAU: Addis Ababa University:
ALT: alanine aminotransferase:
ALP: alkaline phosphatase:
AST: aspartate aminotransferase:
ATDH: Anti-tuberculosis drug-induced hepatotoxicity:
INH: isoniazid:
NGO: Non-governmental organization:
SAM: Swiss Albino Mice:
SGOT: serum glutamate-oxaloacetate transaminase:
SGPT: serum glutamate-pyruvate transferase:
pNPP: p-Nitrophenylphosphate:
pNP: p-Nitrophenol:
TB: Tuberclosis:
TBIL: total bilirubin:
NADH: nicotine amide adenine dinucleotide:

## Acknowledgments

The authors wish to thank Dr. Frank Ashall, Dr.YididiyaBelayneh, TikurAnbessa Specialized Hospital, Addis Ababa University, Aksum University.

---

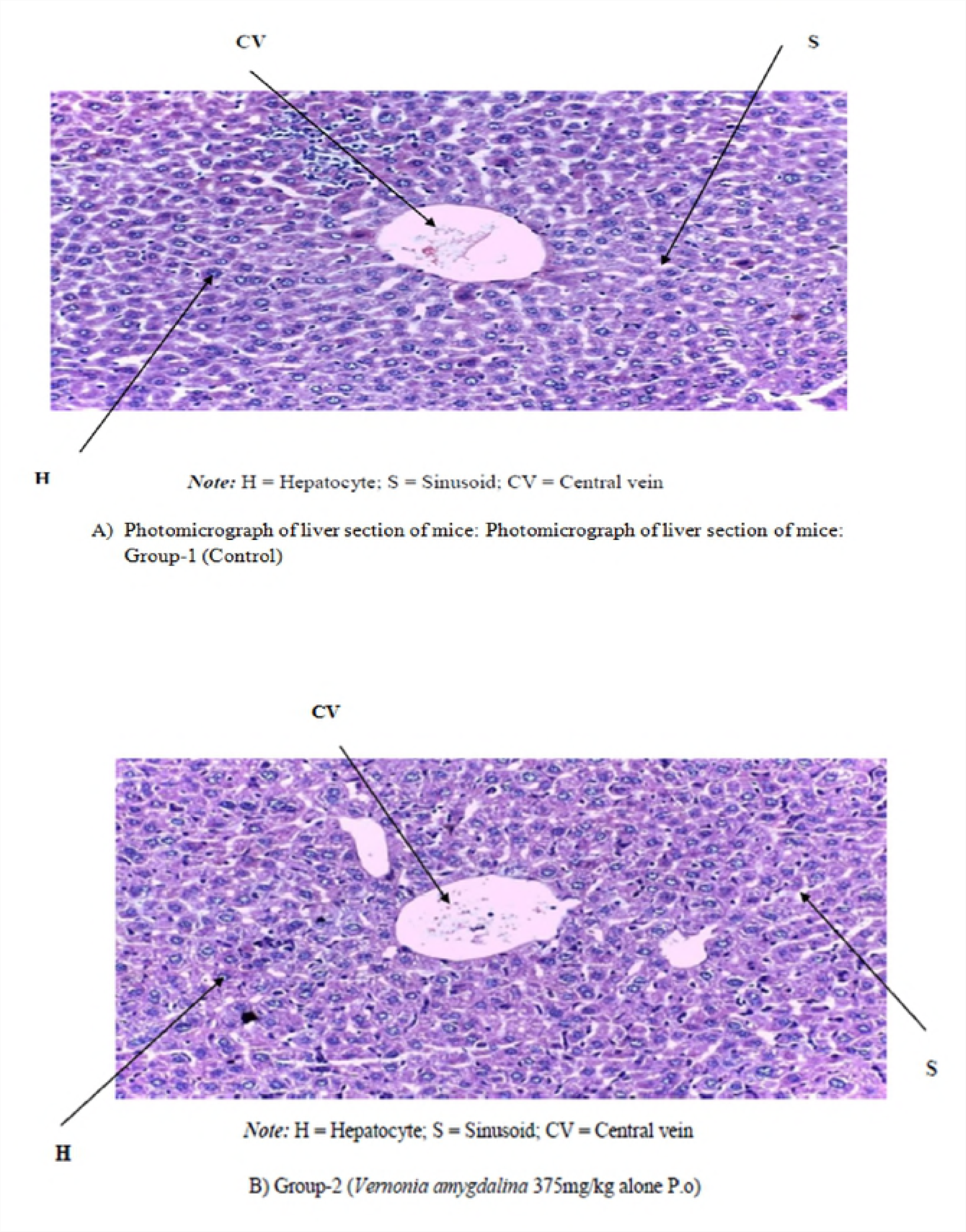

## References

1. Singh A, Bhat TK, Sharma OP. Clinical Biochemistry of Hepatotoxicity. J Clinic Toxicol. 2011; S4(001).

2. Naruse K, Tang W, Makuuchi M. Artificial and bioartificial liver support: A review of perfusion treatment for hepatic failure patients. World J Gastroenterol. 2007; 13:1516–1521.

3. Dhingra M, Nain P, Nain J, Malik M. Hepatotoxicity v/s hepatoprotective agents a pharmacological Review. International Research Journal of Pharmacy. 2011.

4. Saukkonen J, Cohn DL, Jasmer RM, Schenker S, Jereb JA. An Official ATS Statement: Hepatotoxicity of antituberculosis therapy. Am J RespirCrit Care Med. 2006; 174:935–952.

5. Papay JI, Clines D, Rafi R, Yuen N, Britt SD. Drug-induced liverinjury following positive drug rechallenge. Regul Toxicol Pharmacol. 2009; 54:84–90.

6. Deng X, Luyendyk JP, Ganey PE, Roth RA. Inflammatory stress and idiosyncratic hepatotoxicity: Hints from animal models. Pharmacol Rev. 2009; 61:262–282.

7. Vijaya PV, Suja R, Shyamala DCS. Hepatoprotetive effect of Liv.52 on Antitubercular Drug induced Hepatotoxicity in rats. Fitoterapia (LXIX). 1998; 6:620.

8. Mary C, Kasturi S, Parames CS. Herbal protein isolate protects liver from nimesulid induced oxidative stress. Pathophysiol. 2006; 13:95–102.

9. Vuilleumier N, Rossier MF, Chiappe A, Degoumois F, Dayer P, Mermillod B, Nicod L, Jj D, ochstrasser D. CYP2E1 genotype and isoniazid-induced hepatotoxicity in patients treated for latent tuberculosis. Eur J Clin Pharmacol. 2006; 62(6):423–429.

10. Black M, Mitchell JR, Zimmerman HJ, Ishak KG., Epler GR. Isoniazid-associated hepatitis in 114 patients Journal of Gastroenterology. 1995; 69(2):289–302.

11. Mitchell JR, Zimmerman HJ, Ishak KG, Thorgeirsson UP, Timbrell JA, Snodgrass WR, Nelson SD. Isoniazid liver injury: clinical spectrum, pathology, and probable pathogenesis. Ann Intern Med. 1996; 84:181–192.

12. Timbrell JA, Mitchell JR, Snodgrass WR, Nelson SD. Isoniazid hepatoxicity: the relationship between covalent binding and metabolism in vivo.. J Pharmacol Exp Ther. 1980; 213(2):364–369.

13. Arshad AN, Navdeep S, Koyal S., Kale MK. Hepatoprotective Effect of Rimonabant Against Isoniazid Induced Liver Damage in Albino Wistar Rats nternational Journal of Pharmaceutical & Biological Archives. 2010; 11(4):473.

14. Chen et al. he protective effects of ursodeoxycholic acid on isoniazid plus rifampicin induced liver injury in mice. European Journal of Pharmacology. 2011; 659:53–60.

15. CDC. Sever isoniazid-associated liver injuries among persons being treated for latent tuberculosis infection: United States 2004-2008 MMWR. 2010; 59:224–229.

16. Arhoghro EM, Ekpo KE, Anosike EO, Ibeh GO. Effect of Aqueous Extract of Bitter Leaf (Vernonia Amygdalina) on Carbon Tetrachloride (CCl4) Induced Liver Damage in Albino Wistar Rats. European Journal of Scientific Research. 2009; 26(1):115–123.

17. Imir G M, Tetsuya N., Jack U. Direct Oxidation and Covalent Binding of Isoniazid to Rodent Liver and Human Hepatic Microsomes: Humans Are More Like Mice than Rats. University of Toronto, Canada. 2012.

18. Wendy A. Clinical chemistry laboratory perspective. Texas: University of Texas. 2007.

19. Pratap A. Inhibition of endogenous hedgehog signaling protects against acute liver injury after ischemia reperfusion. Pharm Res. 2010; 27:2492–2504.

20. Nyblom H, Berggren U, Balldin J, Olsson R. High AST/ALT ratio may indicate advanced alcoholic liver disease rather than heavy drinking. Alcohol Alcohol. 2004; 39:336–339.

21. Lamberg S., Rothstein R. Laboratory manual of histology and cytology. USA Westport Conn. 1978.

22. Singh D. Principles and techniques in histology micrograph and photomicrography, 2 edn. New Delhi, India: CBS Publishers and Distributors. 2006.

23. Mohan H. Pathology practical book. Ltd, New Delhi. 2007.

24. Clopton R. Harris hematoxylin and eosin-xylol staining protocol: Hotel Intestine Laboratory for Parasitology. 2006.

25. Iwalokun BA, Efedede BU, Alabi-Sofunde JA, Oduala T, Magbagbeola OA, Akinwande AI. Hepatoprotective and Antioxidant Activities of Vernonia amygdalina on Acetaminophen-Induced Hepatic Damage in Mice. Lagos, Nigeria. University of Lagos. 2006.

